# The BELT and phenoSEED platforms: shape and colour phenotyping of seed samples

**DOI:** 10.1101/825695

**Authors:** Keith Halcro, Kaitlin McNabb, Ashley Lockinger, Didier Socquet-Juglard, Kirstin E Bett, Scott D Noble

## Abstract

**Background:** Seed analysis is currently a bottleneck in phenotypic analysis of seeds. Measurements are slow and imprecise with potential for bias to be introduced when gathered manually. New acquisition tools were requested to improve phenotyping efficacy with an emphasis on obtaining colour information.

**Results:** A portable imaging system (BELT) supported by image acquisition and analysis software (phenoSEED) was created for small-seed optical analysis. Lentil (*Lens culinaris* L.) phenotyping was used as the primary test case. Seeds were loaded into the system and all seeds in a sample were automatically and individually imaged to acquire top and side views as they passed through an imaging chamber. A Python analysis script applied a colour calibration and extracted quantifiable traits of seed colour, size and shape. Extraction of lentil seed coat patterning was implemented to further describe the seed coat. The use of this device was forecasted to eliminate operator biases, increase the rate of acquisition of traits, and capture qualitative information about traits that have been historically analyzed by eye.

**Conclusions:** Increased precision and higher rates of data acquisition compared to traditional techniques will help breeders to develop more productive cultivars. The system presented is available as an open-source project for academic and non-commercial use.

## Background

Lentils (*Lens culinaris* L.) are graded and sold based on a variety of visual traits. These differences can be subtle and are influenced by genetic and environmental factors. Researchers evaluate thousands of seed samples by visual inspection and manual dimensional measurements, which are time-consuming and imprecise tasks. There are practical limitations on the number and size of samples that can be assessed which creates limitations in the statistically valid conclusions that can be made. To overcome these restrictions, a complete system was requested by lentil researchers to analyze seeds with the expectation of quantifying size, shape, colour and seed coat patterning. The system would include a machine for image acquisition and software to acquire, process and analyze seed images. Analysis would return quantifiable measurements suited for rigorous statistical analysis to improve the understanding of the genetics of seed quality traits and identify genes relevant to the breeding program.

Devices to enable high-throughput grading of phenotypic traits of seed coats have been explored previously, with some being commercially available. Solutions explored have included flatbed scanners (Shahin and Symons, 2001; Smykalova et al., 2013) and over-head cameras that image many seeds at once (Vibe QM3, Neutec Group Inc., Farmingdale NY, USA; Opticount, Process Vision LLC, Richmond VA, USA; VideometerLab 4, Herlev, Denmark)^[1]^. These methods generally require some method of separating groups of seed physically (SeedCount, Next Instruments, Condell Park NSW Australia) or in post processing. Where multiple seeds are imaged together, the spatial resolution is typically lower compared to the resolution attainable when imaging a single seed. Other commercial systems such as high-speed sorters for commercial seed sorting or those targeted at a laboratory environment (QSorter Explorer, QualySense AG, Glattbrugg, Switzerland) use relatively sophisticated multi-camera imaging systems to image seeds in a particular orientation, and incorporate sorting equipment that may not be required in all settings. Commercial systems provide a convenient, off-the-shelf solution and often bundle in technical support to solve data acquisition issues and provide insight into analysis problems. The ability to adapt commercial solutions to new or established workflows, measurements or analysis will vary with system and vendor.

Software to enable seed phenotyping is often focused on calculating size and shape parameters from 2D images (Tanabata et al., 2012). Expanding into three dimensional shape analysis yields interesting information on seed plumpness and symmetry which are properties associated with highly sought after traits in lentil (LeMasurier et al., 2014; Shahin et al., 2012). To enable calculation of three-dimensional seed shape properties, systems often incorporate several cameras arranged around the staging area to observe a seed from orthogonal positions. With two orthogonal views of the same seed, volumetric properties can be extracted through mathematical means, often with the assumption of modelling the seed as an ellipsoid (Nikam and Kakatkar, 2013; Shahin et al., 2006). Greater refinements include modelling the seed as a tilted ellipsoid by identifying the upper and lower halves, but this necessitates a sharp delineation from the upper and lower surfaces in the image as a result of directional lighting (Shahin et al., 2006). The expansion into identifying ellipsoid tilt increases the precision of height estimations slightly but the lighting requirements are stringent.

Groups have identified the need for accurate colour information when grading seeds and comparing results across devices (Black and Panozzo, 2004; Schettini et al., 1995; Shahin and Symons, 2003) and have worked to develop calibration of images in agriculture while minimizing in-field requirements (Sunoj et al., 2018). Colour calibration brings the colour of any images captured to what would be expected under a standard illuminant (standardized light source definition) to facilitate colour comparisons (Ricauda Aimonino et al., 2013). Colour information can be represented in a variety of ways, including perceptually uniform colour spaces. Perceptually uniform colour spaces such as CIELab (L*a*b*) are preferred for food studies as they represent human perception (Wu and Sun, 2013). Distance between two points in the L*a*b* colour space is directly related to the difference a human could distinguish between the two (Sharma et al., 2005). This provides parity between machine separation of colour by measurements of distance and human separation of colour by perception (Gentallan et al., 2019; Valadez-Blanco et al., 2007).

The requirements presented by researchers interested in seed analysis at the University of Saskatchewan described a means of quantifying seed colour and shape statistics for a large number of samples (>10,000 annually), based on representative sub-samples containing 100 to 200 seeds. Previous practice was a visual assessment of these sub-samples for qualitative colour classification, and size characterization using either standard sieves and/or manual measurements of a small number (<10) of seeds using calipers. A speed target for image acquisition of one 200-seed sample per minute was set, representing a significant increase in throughput and information available over these manual methods.

Workflow was a key design consideration given the large number of samples. Solutions that could be made broadly available and support the development of new measurement and analysis techniques were desirable. This problem sat in a gap between existing solutions; existing inexpensive or open-source approaches did not inherently meet the desired throughput but could have been overhauled with significant work. Commercially available lab products were not found that met all the requirements for workflow, portability, throughput, resolution or image quality, or openness. This led to the development of a hopper-fed, seed-singulating, conveyor-based system for image acquisition (BELT), and a set of batch-processing analysis codes that are scalable for progressively analyzing (and re-analyzing as required) the forecasted terabytes of image data (phenoSEED). A cross section render of BELT (Figure 1) and the phenoSEED flowchart (Figure 2) are presented here to contextualize the remainder of this paper.

**Figure 1.**
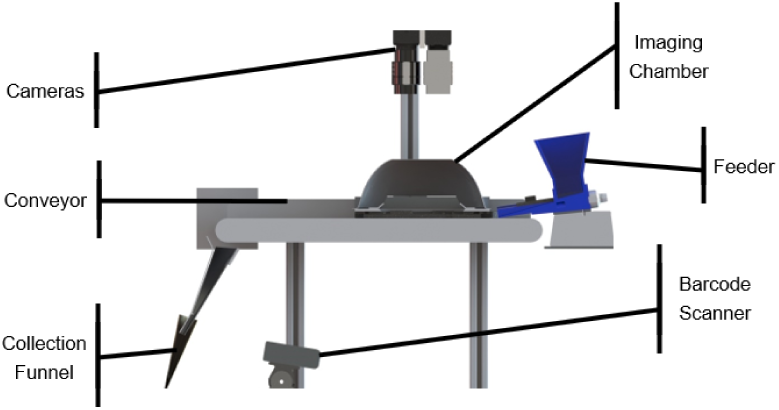
BELT. Cross section render of the physical BELT system.

**Figure 2.**
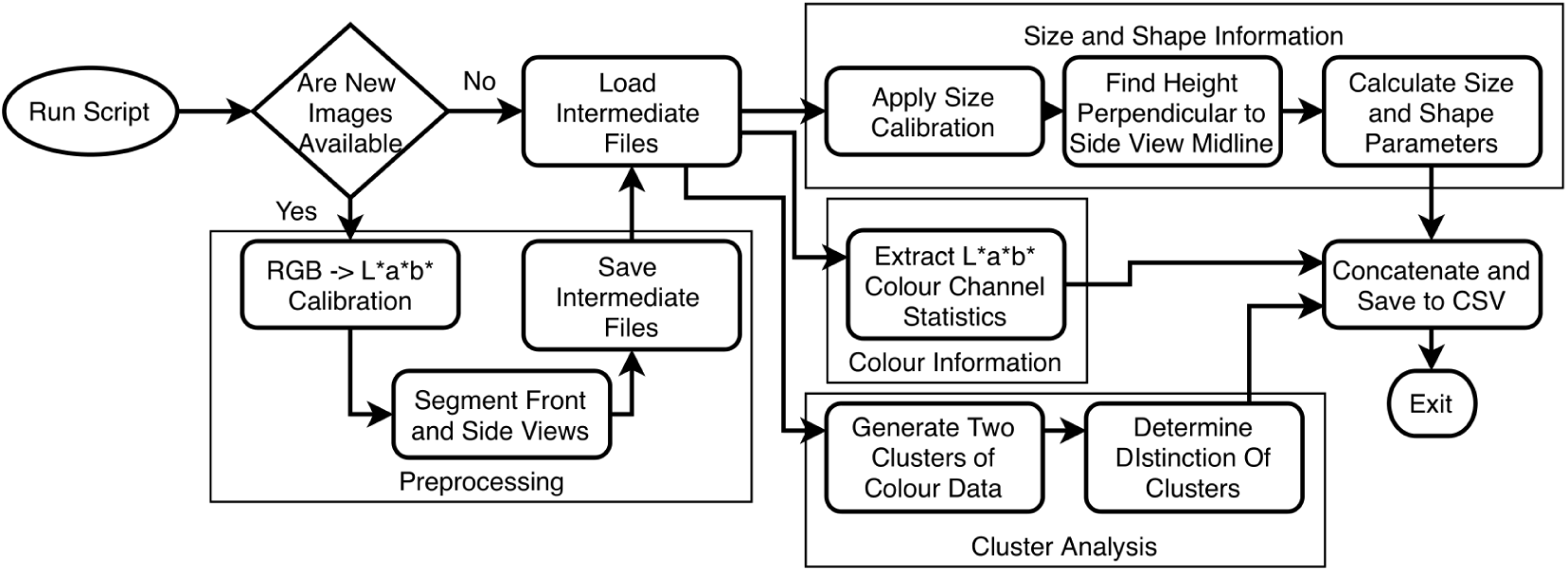
Seed image processing flowchart. Preprocessing is conditional and can be skipped.

## Results

### Image Acquisition Performance

The overall performance of the data acquisition system met the design objectives of speed and ease of acquisition. The acquisition system used a barcode scanner to read the sample envelope labels already in use by the breeding project which prevented errors arising from manual entries of identifying information. Seed images acquired by BELT were immediately displayed on the control GUI to give feedback to the user and allowed for multiple users to be trained easily and quickly. Each seed image contained a top-down view which included a reflective prism to show a side view of a passing seed as shown in Figure 3. Camera settings were locked in and hidden from the operator, enabling both straightforward operation and consistent imaging settings. Consistent settings led operators to trust in expected results and diagnose errors quickly. Static lighting and camera light-gathering parameters allowed colour calibration to be determined once and applied without adjustment over an extended working period.

**Figure 3.**
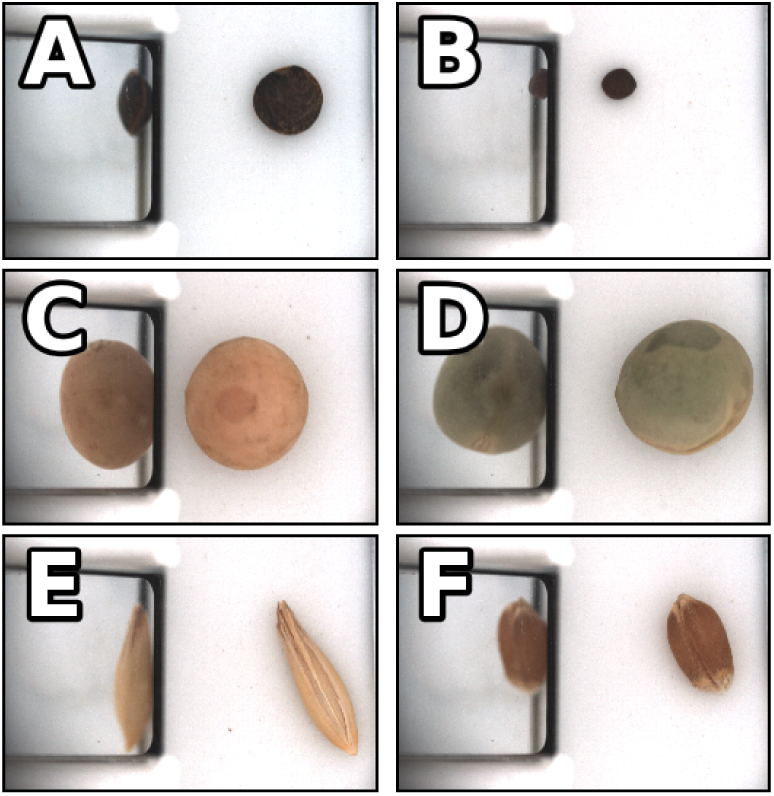
Sample images of several seeds. A) lentil with a dark seed coat B) canola C) yellow pea D) green pea E) oat F) wheat.

The vibratory feeder that carried seeds from the hopper to the conveyor has proved to require some attention from the operator. Due in part to the wide range of physical properties of the samples, flow continuity can vary. The angle of the vibratory feeder was adjustable to make certain that seeds of varying sizes were singulated properly, and an adjustable gate was implemented to help manage flow from the hopper. It has been observed that operators tend to gently agitate the seeds to maintain steady flow from the hopper. While the design has been effective considering its simple construction (a 3D-printed feeder mounted on a generic aquarium air pump), it has been identified as an area for improvement in the design.

Optical triggering was based on 3 mm diameter infrared light emitting diodes and phototransistors that were installed above the conveyor belt. These diodes and phototransistors sat too high off the conveyor belt to ensure reliable triggering by very small seeds (wild-type lentils or canola seeds). These triggers were re-implemented using 1.6 mm surface-mount components. This has also meant that small stones or other debris would cause false triggers resulting in images that do not contain seeds. An area of ongoing work in post-processing is distinguishing faulty images of non-seed items or broken and damaged seeds.

### Size Calibration

A ruler with one-millimeter deviations was imaged under three conditions – with face up sitting on the conveyor, facing the prism while near the prism edge, and facing the prism at a far distance. Images of a ruler in the imaging chamber (Figure 4) were analyzed to create linear fits of pixel counts to physical distances. The marked divisions of the ruler were horizontal in the image, and an edge filter was used to find the vertical position of the edges of the marks. Summing the edge filter along rows created a periodic series that had peaks associated with transitions between white and black areas. The ruler divisions were plotted against the pixel position of positive peaks with a simple linear fit. The processing steps for size calibration are shown in Figure 5. The results of the camera calibrations are presented in Table 1. Any conversion of size or shape information extracted from the top view to physical measurements applied the size calibration directly. Any side view items like height linearly interpolated a value of size calibration based on the horizontal centre position seed as a proxy for distance to camera through the mirrored prism.

**Table 1.** Information of linear regression fits of camera pixel size. Each cell has two entries, one for each camera. R^2^ values for each fit line were above 0.99. Note that the side view scaling is linearly interpolated between near and far view based on the midpoint of the object relative to the end points.

| View | Camera | Slope [mm/px] | Intercept [mm] | End Point [px] |
| --- | --- | --- | --- | --- |
| Top | A | 0.0094 | 0.012 | - |
|  | B | 0.0095 | -0.006 | - |
| Near Side | A | 0.0093 | 0.002 | 1000 |
|  | B | 0.0094 | 0.009 | 1250 |
| Far Side | A | 0.0101 | 0.008 | 2200 |
|  | B | 0.0102 | -0.016 | 200 |

**Figure 4.**
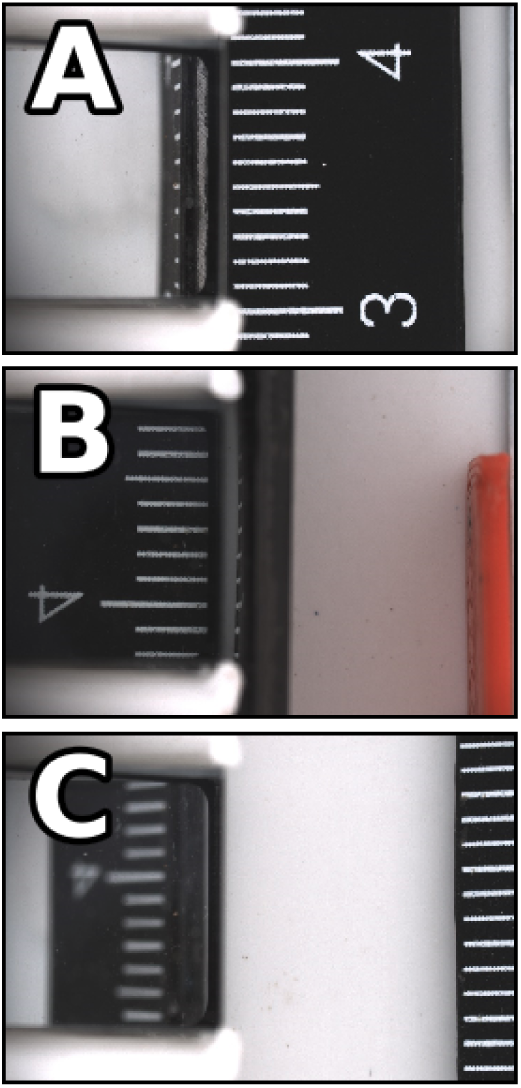
Sample height calibration images. A) a top view B,C) two side views with different distances from the prism. Note the depth of focus is shallow and the side view of the ruler when it is far from the prism is not nearly as sharp as the other views.

**Figure 5.**
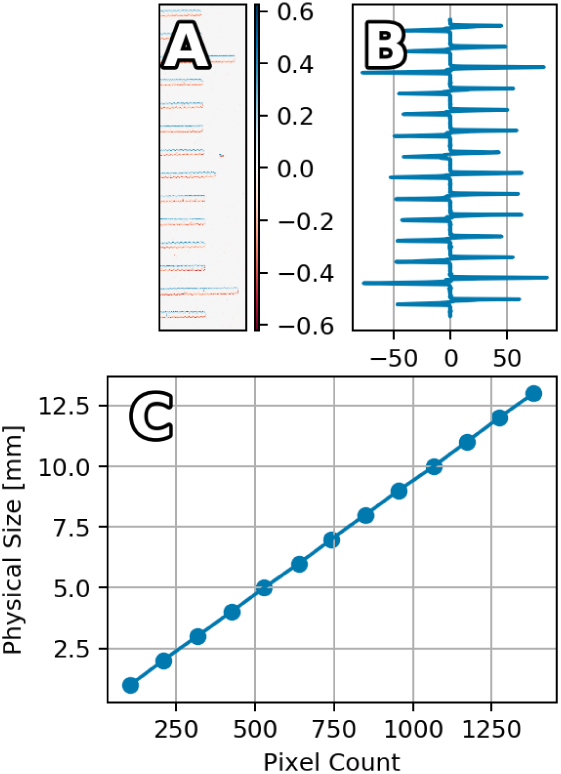
Height calibration process. A) a cropped image of a ruler after application of the sobel filter B) the results of summing the sobel filter image along rows C) positive peaks in the signal are plotted against the distance between ruler tick marks.

#### Cross-camera Validation

The calibration values of each camera were very close to the other, which was expected as they had the same lens and sensor combination and were mounted at similar heights above the target. To test the cross-camera compatibility of the calibrations, circular targets cut from plywood with a nominal diameter of six millimetres were imaged with both cameras. The diameter of each target was measured with a caliper five times then was averaged, while the mean equivalent diameter of the circular targets was extracted from three images acquired with BELT and segmented with phenoSEED. Diameter measurements of the circular targets are contained in Table 2. The extracted values were overestimated by less than one percent on average, but the measurements from each camera were very closely related with 0.21% difference between them on average.

**Table 2.** Summary information of size validation between two cameras. The target diameter was measured with calipers and was also extracted from images from each camera. Percentage differences were calculated between the image and target diameter as well as between both image diameters.

| Target Diameter [mm] | Camera | Image Diameter [mm] | Difference | Intra-camera Difference |
| --- | --- | --- | --- | --- |
| 5.87 | A | 5.89 | 0.34% | 0.20% |
|  | B | 5.91 | 0.68% |  |
| 5.98 | A | 6.02 | 0.67% | 0.16% |
|  | B | 6.03 | 0.84% |  |
| 6.09 | A | 6.09 | 0.00% | 0.26% |
|  | B | 6.10 | 0.16% |  |

### Colour Calibration

The transformation between RGB image values to calibrated L*a*b* values was assumed to be non-linear and could be described by an artificial neural network. A Multi-layer Perceptron for Regression (MLPR) was initialized with default values of the implementation in scikit-learn (one layer of 100 hidden neurons) (Pedregosa et al., 2011). No work was done on investigating parameters that fine-tune the learning algorithm such as adding additional hidden layers, changing the learning rate or adjusting the decay of moment vectors in the Adam solver.

The results of the non-linear colour calibration fits are displayed in Figure 6, in which the values of each L*a*b* colour channel before and after calibration are plotted against the supplied values for the X-Rite ColorChecker Digital SG (X-Rite, Grand Rapids, MI). In both cameras, the uncalibrated lightness values were offset slightly but the fits were poor as noted by the RMSE reported in Table 3. The calibration aligns the values per channel very closely to the expected values as noted by slopes close to one and high coefficients of determination reported in Table 4. The colour intensities of the uncalibrated data was very low, as the slope values show the response to colour information was only sixty percent of what would be expected. Calibration is most effective in the colour information, as can be visually perceived by comparing the lentil image in Figure 3 to the same lentil image after calibration is applied which can be viewed in Figure 7.

**Table 3.** Comparisons of linear fits between measured L*a*b* images of an X-Rite ColourChecker Digital SG to the ground truth L*a*b* values. Each cell has two entries, one for each camera.

| Colour Channel | Camera | Measured Data |  |  |
| --- | --- | --- | --- | --- |
| | | Slope | $R^2$ | RMSE |
| $L^*$ | A | 0.954 | 0.894 | 13.6 |
|  | B | 0.997 | 0.895 | 12.1 |
| $a^*$ | A | 0.537 | 0.872 | 13.1 |
|  | B | 0.584 | 0.871 | 12.2 |
| $b^*$ | A | 0.551 | 0.865 | 15.6 |
|  | B | 0.594 | 0.871 | 14.6 |

**Table 4.** Comparisons of linear fits between calibrated L*a*b* images of an X-Rite ColourChecker Digital SG to the ground truth L*a*b* values. Each cell has two entries, one for each camera.

| Colour Channel | Camera | Calibrated Data |  |  |
| --- | --- | --- | --- | --- |
| | | Slope | $R^2$ | RMSE |
| $L^*$ | A | 0.991 | 0.991 | 2.10 |
|  | B | 0.986 | 0.990 | 2.71 |
| $a^*$ | A | 0.982 | 0.986 | 2.77 |
|  | B | 0.991 | 0.983 | 3.26 |
| $b^*$ | A | 0.994 | 0.993 | 2.90 |
|  | B | 0.994 | 0.990 | 3.00 |

**Figure 6.**
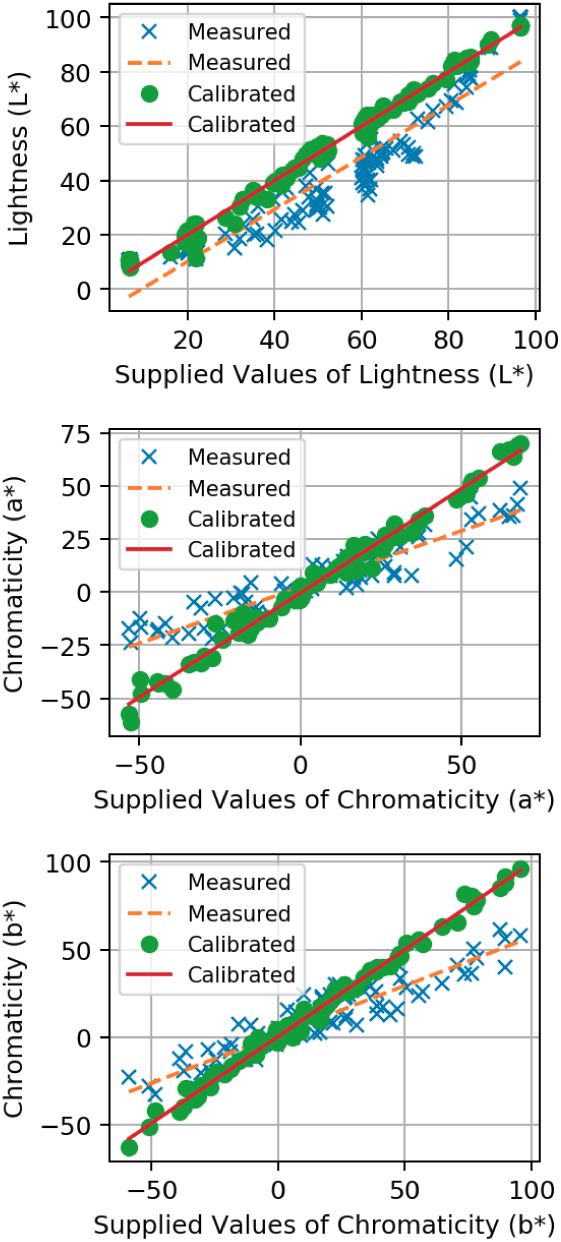
Colour calibration in the L*a*b* colour space. Plotted relationships between measured and calibrated data to the supplied L*a*b* values for a subset of a ColorChecker Digital SG.

**Figure 7.**
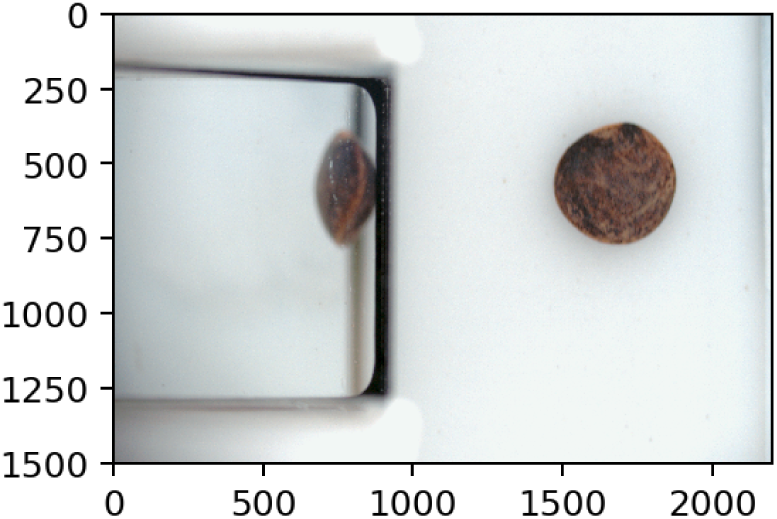
A sample lentil with a dark seed coat after colour calibration. This is the same lentil image showcased in Figure 3.

#### Cross-camera Validation

The differences between the predicted values and calibrated values were calculated using two standards for colour difference (Sharma et al., 2005) implemented in scikit-image (van der Walt et al., 2014). Colour differences between the supplied L*a*b* values and the measured values before and after calibration are presented in Table 5 with the old standard of ΔE* CIE76 and the newer ΔE* CIEDE2000 (Sharma et al., 2005). Calibration reduced the colour distances significantly (Table 5) and it is worth re-iterating that the colour difference between L*a*b* points is proportional to the degree of distinction in human perception. Under a just noticeable difference (JND) threshold (ΔE ≈ 2.3) the colour values can be considered to be perceptually the same (Mahy et al., 1994). The visual distinction between the original PNG images and the calibration dataset was roughly four times greater than the visual distinction between calibrated images and the dataset which was just above the JND threshold. When comparing the predicted datasets against each other, the mean ΔE* CIEDE2000 distance was 3.23 (s.d. 1.9) while the mean ΔE* CIEDE2000 distance between the original images of the dataset was 2.69 (s.d. 1.9). The calibrated images from different cameras were on average as distinct from each other as they were independently distinct from the training set, and applying the non-linear calibration caused the images from each camera to diverge slightly from each other. Samples that diverged significantly between predicted and actual values (ΔE* CIEDE2000>5) were usually dark (L*≤20), which suggested that the training process struggled in fitting those samples. This can be viewed in Figure 6 where a cluster of several calibrated points with L* values equal to twenty were below the fit line, meaning those calibrated datapoints were darker than the actual values.

**Table 5.** Colour distances between measured and calibrated data to the ground truth L*a*b* values. Each cell has two entries, one for each camera.

| Dataset | Camera | $\Delta E^*$ CIE76 | $\Delta E^*$ CIEDE2000 |
| --- | --- | --- | --- |
| Measured points | A | 21.1 (s.d. 12.4) | 13.3 (s.d. 6.47) |
|  | B | 19.2 (s.d. 11.8) | 12.0 (s.d. 6.40) |
| Calibrated points | A | 3.80 (s.d. 2.76) | 2.59 (s.d. 1.71) |
|  | B | 4.45 (s.d. 2.56) | 3.01 (s.d. 1.74) |

#### Clustering

K-means clustering and Gaussian Mixture Models were investigated for clustering colour data. Both algorithms were instructed to find two clusters, with the expectation that one would represent a base colour and the other would represent a pattern colour. K-means clustering randomly assigns points as cluster centres, then all points are assigned to a cluster based on proximity and the cluster centres are updated to be equal to the cluster mean. This iterative process continues until cluster definitions stop changing. A Gaussian mixture model attempts to find and describe a given number of Gaussian distributions in the dataset and then calculates the maximum likelihood to assign each sample to a cluster. The average log-likelihood of the GMM can be returned when a prediction is made using a trained model. The application of K-means and GMM clustering on three sample lentils (two single-colour (A and C) and one patterned (B)) is shown in Figure 8.

**Figure 8.**
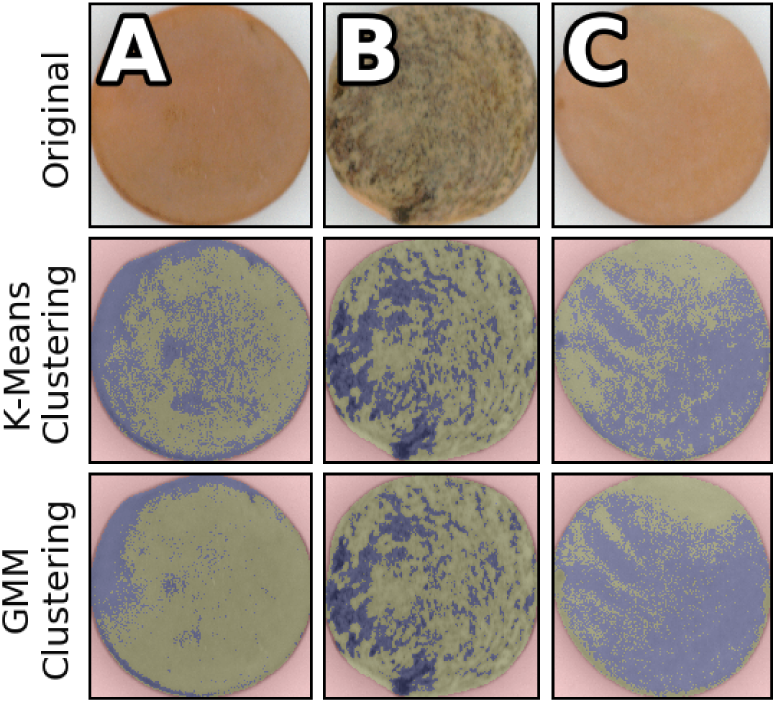
The results of clustering lentil seed colour data. Clusters are highlighted in blue and green. Background pixels are red. Row 1) original lentil images cropped to the bounding boxes Row 2) the results of K-means clustering Row 3) the results of Gaussian mixture model clustering.

Each clustering algorithm was looking for two colours and their performance on single colour lentils differed significantly. K-means clustering was ultimately not appropriate for this application as the algorithm would attempt to maximize the distance between the two equally sized clusters when applied to a single colour lentil as noted by the cluster populations in Table 6. K-means and GMM had similar performance on patterned lentils with as they both report color distance of about ten (ΔE* CIEDE2000≈10). The colour distance between cluster centres for examples A and C are below JND. Single-colour lentils could be distinguished by their small distance between cluster centres and a less negative average log-likelihood as distributions overlapped and samples were assigned with less certainty. Running an algorithm looking for two clusters was necessary to find seed coat patterns but the GMM allowed for single-colour lentils to be identified based on the scoring of the fit.

**Table 6.** Summary of Euclidean distances between clusters, self-reported scores and cluster populations when L*a*b* colour pixels are clustered. Example labels refer to Figure 8. The cluster distances were calculated via ΔE* CIEDE2000.GMM: Gaussian mixture model

| Clustering Metric | Example A | Example B | Example C |
| --- | --- | --- | --- |
| K-means Cluster Distance | 2.82 | 10.9 | 2.47 |
| GMM Cluster Distance | 2.25 | 9.20 | 1.50 |
| GMM Average Log-likelihood | -6.57 | -8.04 | -6.09 |
| K-means Cluster Population (Green/Blue) | 62% / 38% | 70% / 30% | 38% / 62% |
| GMM Cluster Population (Green/Blue) | 82% / 18% | 72% / 28% | 32% / 68% |

#### Sample Study

To assess the use of BELT and phenoSEED in creating useful information, a group of ten lentil samples was assessed to find the distribution of size, mean seed coat colour and shape descriptors (Equations 1, 2 3) in each sample. Boxplots of the distribution of the parameters of three samples of wild (W) lentil varieties and seven samples of cultivated (C) lentils are shown in Figure 9. The wild lentils were generally smaller, darker and more spherical than the cultivated samples. Some samples tended to have many outliers in all of the measurements, suggesting that the images with seeds were not segmented correctly or images without seeds were passed to phenoSEED. Automatic outlier removal was not implemented in phenoSEED as broken or otherwise abnormal lentils are potentially valuable as sources of phenotypic information related to handling characteristics.

**Figure 9.**
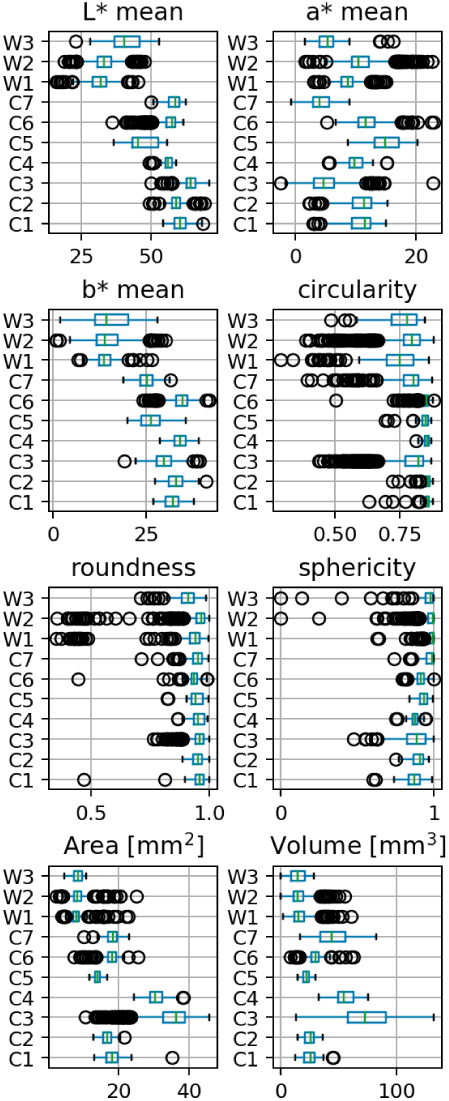
phenoSEED information for wild and cultivated lentils. Boxplots of shape, size and colour information for cultivated (C) and wild (W) lentil varieties.

## Discussion

### Data Processing Rates

Over the course of an eight hour workday, BELT could gather up to 200GB of image data as the system was fast, automated, and required minimal operator oversight. This is a significant amount of image data to move and it took several hours to upload to network storage as the average file movement speed on the network was 80 MB/s. The phenoSEED Python script spawned 16 workers to process input files in parallel using a multiprocessing pool. The implementation of a multiprocessing pool makes it very difficult to profile the code without extensive changes in the programming style. There is no detailed function by function breakdown of time, but the program does a naïve timing of the entire analysis then divides by the number of files to get an overall average of time per one process. The script was able to process a 9.45 MB input PNG image in 1.35 seconds on average. This resulted in a processing rate of 7 MB/s, and roughly eight hours of processing time for an input of 200GB.

phenoSEED included a preprocessing step that would transform image data and discard extraneous information. The results of preprocessing are termed intermediate files and when working from the intermediate files and skipping preprocessing, the script was able to extract shape, size, colour and clustering information in 0.35 seconds per intermediate file. Clustering is the major time sink; without it all other stats were extracted at a rate of one intermediate file per 0.065 seconds.

### Future Development

phenoSEED separated preprocessing and main processing while also splitting main processing into several distinct functions that are clearly invoked by the top-level function. Existing functions could be easily modified, or new ones added, with minor revisions. Any revised script can be deployed to analyze the files preprocessed by a previous run of the program. Size, shape and colour information are common elements of seed selection, and any of these general traits could be combined with additional seed-specific information similar to how seed coat patterning analysis was developed for lentils. Highest among suggested improvements when analyzing lentils would be to extract information about the morphology of the colour clusters to feed into a classifier to classify seed coat patterns. Investigating a smaller sample size to fit the GMM colour clustering algorithm would improve overall processing speed significantly, but work is required to validate the results with different sizes of training populations. Python packages used in phenoSEED are all open-source which allow for redistribution. The analysis script currently cannot be entirely divorced from the acquisition system. The main processing functions can be used as a starting point for other seed image analysis projects.

BELT can acquire images of many types of seeds and phenoSEED works with files created by BELT by parsing the known file structure. Refinement to the vibratory feeder design is being considered to effectively handle more seed types. Currently there is a gate at the hopper to control the opening to the channels and the angle of descent can be adjusted. Flat disk-shaped seeds like lentils tended to singulate well while larger rounder seeds like peas would roll down the feeder and pass the imaging chamber while moving faster than the belt speed. An end-gate at the interface between the channel and the belt would help to queue the larger seeds. The image acquisition subsystem was extremely sensitive to ensure seeds of varied sizes would all be captured correctly. The LabView acquisition program that handles camera initialization and triggering can be bundled into an executable and distributed.

## Conclusions

A high-throughput phenotyping system was developed to easily and quickly score visual information of lentil seeds. A conveyor system was assembled on a portable cart that held two cameras and associated LabView imaging software to create BELT. BELT automated seed movement through an imaging chamber which provided a good environment to capture images. An operator had to load and unload packets of seeds but did not influence the acquisition process. LabView software enabled easy initialization and automatic image acquisition with a user-friendly touchscreen GUI to allow users to check images as they were captured. Complete automation of data transfer was not implemented, and operators were responsible to push their results to a network daily.

Image analysis was supported by scripts that automated data transfer, while phenoSEED processed images and returned results of extracted shape, size and colour. The phenoSEED python script was written with modularity in mind, so that new inquiries and functions can be built and re-deployed quickly on pre-processed data. Preprocessing reduced the data volume stored by eighty percent and reduced the time to analyze by seventy percent during subsequent runs. Once the intermediate files had been generated correctly with accurate segmentation and colour correction, preprocessing was no longer required when running the script again with new functions. Preprocessing reduced disparities between the images and their calibration targets but did not completely align colour information from both cameras. After calibration, the mean colour difference between cameras is roughly the same as the mean colour difference between each camera and the calibration dataset. Preprocessing did not completely decouple the camera from the images as the size scaling factor was still required in the main processing functions.

Development of the analysis script could lead to extraction of better descriptors of lentil seed coat traits, particularly patterning density and pattern type. Analysis of other seeds using this imaging platform should be able to use preprocessing functions to isolate seeds in their images, while main processing functions can be used as they are or adjusted slightly to extract information more useful to the researchers working with that plant.

We are in the process of making the code and hardware plans available for academic and non-commercial use. It is hoped that this will support the collection of more easily cross-comparable data and encourage other research groups to contribute to further development of the project. At the time of publication, a version of the processing script is available at https://gitlab.com/usask-speclab/phenoseed. Further information on a comprehensive hardware and software bundle will be made available as it is packaged for distribution.

## Methods

### BELT System Design and Description

BELT (Figure 1) was designed around a 150 mm wide conveyor with at white, low-gloss belt (Mini-Mover Conveyors, Volcano CA) mounted on an audio-visual cart for ease of mobility between labs. The cart held a small PC with a Windows operating system running a touchscreen display. A small vibratory feeder developed in-house was mounted on the right-hand end of the belt. The operator empties a sample envelope into the feeder, scans the barcode identifying the sample on the envelope, and places the empty envelope under the collection funnel at the left-hand end of the belt. After verifying the barcode was scanned correctly, the operator begins the scan via the touchscreen interface. This starts the conveyor and feeder, forming two lines of singulated seeds on the conveyor belt. Splitting the sample was done to increase the throughput of the system without adding significant complexity. The number of parallel pathways possible is primarily a function of belt and lighting chamber size, number of cameras used, and bandwidth available on the computer.

Once on the conveyor, the singulated seeds move into the imaging dome. The dome interior is painted with flat white paint to diffuse light and minimize shadow. The dome was illuminated with 6500K colour temperature, high colour-rendering index LED strips attached to raised surfaces and pointed towards the interior walls of the dome to provide diffuse lighting conditions (ABSOLUTE SERIES D65, Waveform Lighting, San Francisco CA).

Two 5MP RGB cameras (Chameleon3 CM3-U3-50S5C-C5 USB, FLIR Machine Vision, Wilsonville OR) with 35mm machine vision lenses (LM35JC5M2, Kowa American Corporation, Torrance CA) and extension tubes for magnification were installed above two holes in the top of the imaging chamber. The cameras were triggered by infrared emitter-detector pairs contained in the rails separating and containing the lines of seeds passing through the dome. The central dividing rail also contained right-angled prisms, positioned to provide a partial side view of the passing seed to the camera. The side view reflected in the prism was intended to provide information on shape and height of the lentils without additional cameras. The triggering system worked for seeds of other sizes, as demonstrated by the images shown in Figure 3 which were captured for canola (an oilseed), lentil, yellow and green pea (pulses) and oat and wheat seeds (cereals).

LabView (National Instruments, Austin TX) handled initialization of the USB-cameras. Digital gain for both cameras was set at zero. Exposure time, lens aperture, and belt speed were adjusted to obtain an acceptable balance between image brightness, depth of field and motion blur. Focusing priority was given to the top-down rather than side view. White balance per camera was set once through NI Max (National Instruments) then locked in to avoid the white balance auto-adjusting. Captured images were output to the display for an operator to perform checks. Images were saved locally, sorted into a folder structure based on sample ID, camera, and image number.

### Data

Each camera created a 2200×1500 pixel 24-bit RGB PNG (portable network graphics) image that required 9.45 MB of storage space. Images were saved on a local drive before being moved to a storage server manually at the end of the workday. A Linux PC used a scheduled script to fetch images from the storage server using the Globus file transfer protocol (www.globus.org), call the Python processing script, and then push results back to the storage server. Preprocessing steps reduced data volume significantly by cropping images to the seed and discarding extra information.

### Image Analysis (phenoSEED)

A single Python script (Python Software Foundation, www.python.org) was developed to analyze all seed images. To mirror traditional methods, quantitative traits were extracted including size, shape and colour distribution of the seed coats. Qualitative traits such as patterning type and patterning intensity were secondary goals in the image processing. With the expectation that terabytes of data would have to be processed throughout the project, data handling and processing speed were important considerations. Analysis was required to perform well on all images in the dataset, which included several different colours, shapes and sizes of lentil seeds.

The Linux PC used that ran phenoSEED was equipped with an Intel E5-2660 8-core @ 2.2GHz CPU and 64GB of DDR3 RAM. The Python script was a single file with a main function that spawned 16 processes using Python’s multiprocessing package. Each process ran a top-level function with a reference to a single image file that called all necessary functions to load and pre-process the supplied image and save the intermediate files. A flowchart of the script is shown in Figure 2.

#### Preprocessing

Preprocessing applied a colour calibration to the images, segmented images to extract the seed regions of interest, and output the pre-processing results. Segmentation of seed images relied on accurate colour information to isolate the target seed from the background. Accurate segmentation of the target seed created a one-bit depth image where pixels with a value of 0 were considered background and pixels with a value of 1 represented the shape of the target. The one-bit image mask could be overlaid on the colour image to extract colour information of just the target seed area.

#### Colour Calibration

Datasets for colour calibration transformations were created by imaging 117 squares (one greyscale row and one greyscale column were omitted) of the X-Rite ColorChecker Digital SG for each camera as 24-bit RGB images. The ColorChecker Digital SG contains 140 colour swatches that span a wide colour gamut. Compared to the regular X-Rite ColorChecker, the Digital SG has an increased focus on natural tones of browns and greens which provides more calibration data in the range of expected lentil colours. Forty RGB pixels were selected from each calibration image and were scaled to floating point values by dividing by 255, the maximum value of an unsigned 8-bit integer. Each scaled RGB datapoint was associated with the supplied L*a*b* colour value, setting up a simple dataset of 4680 samples. A Multilayer Perceptron for Regression (MLPR) was used to fit a transfer function from captured RGB to supplied L*a*b* values for each camera as non-linear mapping has shown greater accuracy than linear or quadratic mapping (León et al., 2006; Schettini et al., 1995). An MLPR is a supervised learning neural network implemented in the scikit-learn Python package that excels at modeling non-linear functions (Pedregosa et al., 2011). Ninety percent of the RGB data were used for training and ten percent of the data were kept to validate the model performance during training. One MLPR per camera was initialized with a single hidden layer of 100 neurons using rectified linear unit (ReLU) activation to calculate L*a*b* values. The network feedback and learning was handled by an Adam solver (Kingma and Ba, 2014) which updated neuron weights to minimize the mean squared error between the expected and calculated L*a*b* values. Several measurements of each square of the calibration target were needed as Adam optimization was based on stochastic gradient descent methods and was designed to work with large databases (Kingma and Ba, 2014) and requires more data than quasi-newton methods of fitting a non-linear transfer function (Pedregosa et al., 2011). The application of the MLPR to a lentil seed image can be viewed in Figure 7.

#### Segmentation

Each RGB seed image was colour corrected and transferred to the L*a*b* colour space by the camera-specific MLPR. The L*a*b* image was cropped into a top view (limited to the conveyor) and a side view (limited to the reflection in the prism). Both colour channels (a* and b*) had very small values for the white or black background so the absolute values of each were individually scaled from zero to one then added together to create merged colour channel images, one for the top view and one for the side view. The lightness channel was high for the white backgrounds in the top view and side view except for a vertical band in the prism reflection. The lightness channel was scaled from zero to one then multiplied by a scaling factor of 0.5 before it was subtracted from the merged colour image to create a final merged image. The final merged images had high values in the region of the seed which was surrounded by low value background areas. The mask was primarily based on colour information, but the addition of lightness information allowed for segmentation of dark seeds while scaling factors reduced the effect of dark spots on the belt or reflected in the prism. Each merged image was thresholded using Otsu’s method (van der Walt et al., 2014) to create an image mask of the top or side area of the seed. After filling holes in the objects, the largest remaining object was used as the one-bit image mask of the seed. The steps of segmenting a sample lentil image are shown in Figure 10. Note that the lentil image in Figure 10 has been recombined to the full image from the split views for the purpose of illustration. Any mask that touched the border of the side or top view was automatically excluded from processing to avoid calculating size and shape data for items that did not have a clear side view.

**Figure 10.**
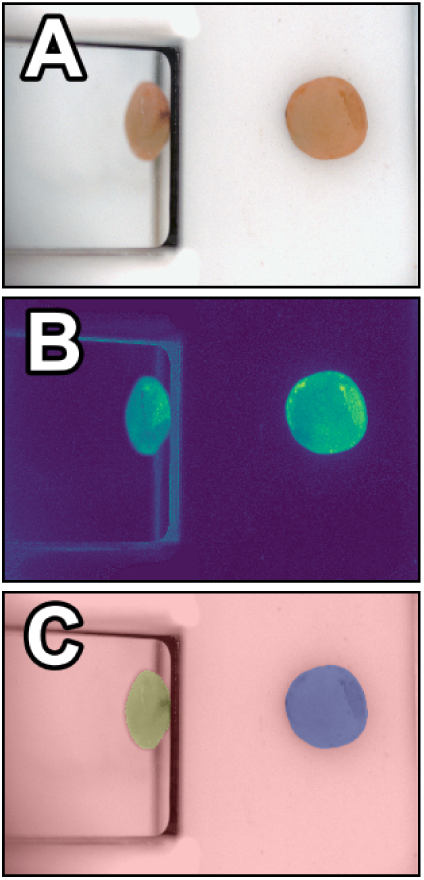
Steps of segmenting the views of a lentil. A) original image B) a merged image of the scaled values of a*, b* and -L*/2 added together C) results of segmentation, with the top mask overlaid in blue and side mask overlaid in yellow. Not shown are the intermediate steps of cropping to the top and side views.

#### Preprocessing Output

The horizontal centre of the top-down segmented view was saved as a marker of the lentil position in the coordinates of the original image. Four items were saved as intermediate files: a colour calibrated cropped L*a*b* image, a top view mask, a side view mask and the horizontal position of the centre of the top view mask. The intermediate files were intended to be stored and utilized when needed to calculate information about lentil seed coats. Regenerating the intermediate files was costly in terms of computation time but is only necessary if colour calibration or segmentation are altered.

#### Main Processing

Following the pre-processing steps, the main processing step extracted morphological and colour statistics from the data and applied a clustering algorithm to the colour data. The one-bit depth image masks were used as sources of shape information. The major axis length, minor axis length, area and perimeter of the top view were calculated in units of pixels. The height was calculated by doubling the length of the longest perpendicular line segment from the midline of the side view to the top edge of the side mask. These parameters are overlaid on an image in Figure 11. All methods that calculate size and shape parameters report their results in pixel count and were multiplied by a size calibration of millimeters per pixel to acquire physical measurements. Pixel size in the top view was assumed to be uniform as the cameras had narrow field of view and images were centred to avoid lens distortion. The scaling of an object reflected in the prism depended on the distance between the object and the prism which is represented by the horizontal position of the top view calculated during preprocessing. Objects further from the mirror appeared smaller in the reflection and the scaling factor of reflected objects was dependent on the position of the object on the belt.

**Figure 11.**
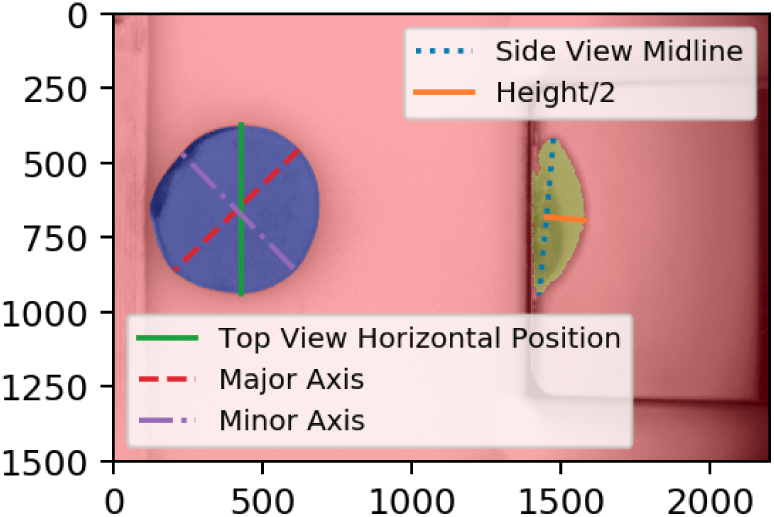
Annotated image showing the locations of several shape parameters. The top mask is highlighted in blue and the side mask is overlaid in yellow.

#### Shape and Size

Roundness and circularity were calculated to describe the 2D shape of the top view, and sphericity described the 3D shape. Roundness and circularity definitions were adopted from those used in the ImageJ software (Schneider et al., 2012). Roundness is the ratio between the area of the shape to the area of the circumscribed circle. Circularity is a ratio of area to perimeter, normalized to a geometrically perfect circle. Sphericity is a ratio of volume to surface area, normalized to a perfect sphere (Wadell, 1935). The surface area and volume calculation approximated the lentil seed as a tri-axial ellipsoid with the 2D major and minor axes and the height as the three axes. All three indices range from 0-1 with 1 representing a perfect circle or sphere.

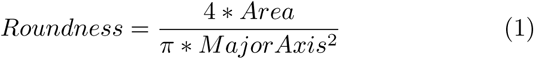

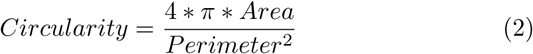

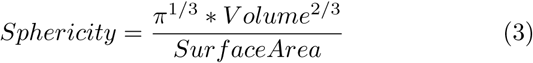

#### Colour Statistics

Descriptive statistics of the colour data over the lentil area were extracted to describe the colour distribution per lentil so that outliers within a sample group could be identified. The statistics extracted were the mean, maximum, minimum and standard deviation of each L*a*b* colour channel. These helped to catalogue the distribution of colours within a sample of 200 seeds.

#### Clustering

The L*a*b* colour values of the top view of the lentil were clustered into two groups using a Gaussian mixture model (GMM), an unsupervised clustering algorithm implemented in scikit-learn (Pedregosa et al., 2011). One GMM per image was fit to ten percent of the available pixels in the top view of the lentil, which was on average 20000 sample points. The model was trained to find two groups within the dataset and was then used to predict the grouping of the rest of the individual pixels in the lentil top-down area. A GMM predicts the grouping of new samples based on maximum likelihood to be in the distributions learned during training. These groups would be assessed to determine seed hull patterning or damage.

## Declarations

### Ethics approval and consent to participate

Not applicable.

### Consent for publication

Not applicable.

### Availability of data and materials

The image datasets captured by BELT used and/or analysed during the current study are available from the corresponding author on reasonable request.

### Author’s contributions

KH developed phenoSEED and calibrated BELT and was a major contributor to the manuscript. KM, AL and SN designed BELT LabView and physical systems. DS and KB provided the impetus for BELT and oversaw operation. DS, KB and SN reviewed the manuscript. All authors read and approved the final manuscript.

## Acknowledgements

The authors would like to thank staff at the College of Agriculture and Bioresources who collected seed samples and operated BELT. Their feedback aided the development of phenoSEED.

## Footnotes

[1] Product and company names are acknowledged as examples of the state of the art or to provide relevant technical detail. This is not intended to be an exhaustive list, nor as an endorsement or criticism of said product or company by the authors.

